# Establishment of a novel method for the production of chimeric mouse embryos using oil droplets

**DOI:** 10.1101/870824

**Authors:** Hiroyuki Imai, Soichiro Tsuda, Tokuko Iwamori, Etsuro Ono

## Abstract

The production of chimeric animals is frequently necessary for the constructing genetically modified animals, and has gained popularity in regenerative medicine in the recent years for the reconstruction of xenogeneic organs. The aggregation method and the injection method are generally used for producing chimeric mice. In the aggregation method, the chimeras are produced by co-culturing embryos and stem cells, and keeping them physically adhered. In the injection method, the chimeras are produced by injecting stem cells into the zona pellucida using microcapillaries. These methods only focus on the generation of chimeric animals, and are not expected to produce reproducible results or allow quantitative evaluation.

This study aimed to establish a novel method for producing chimeric embryos via droplets for improving on the conventional methods that are used for producing chimeric embryos. In this study, the embryonic stem cells and embryos were successfully isolated in the droplets, and the emergence of chimeric embryos was confirmed by co-culture for 6 hours. By this method, the control and operability of stem cell numbers can be regulated, and the method allows better reproducibility and quantification during the production of chimeric embryos. In addition to the conventional methods for producing chimeric embryo, the novel method described herein could be employed for the efficient production of chimeric animals.

## Introduction

The production of chimeric animals is often necessary for the generation of genetically modified animals, and has gained popularity in the recent years for the reconstruction of xenogeneic organs (1,2). There are two classical methods for producing chimeric animals, especially chimeric mice, using pluripotent stem cells, namely, the injection method (3) and the aggregation method (4). The culture conditions for stem cells have been improved for the production of chimeric mice, including the use of chemically defined media or 3i culture system (5, 6, 7). Previous studies have also focused on the number of passages of stem cell lines and the number and ploidy of the host embryos for improving the methods used to induce chimerism (8, 9). These studies have helped improve the production efficiency of transgenic mice that are produced from chimeric mice. However, there are certain issues with the methods that are currently used for producing chimeric embryos, namely, the injection method and the aggregation method, which are discussed hereafter.

Firstly, in the injection method, chimerism is induced by injecting stem cells into the zona pellucida using an microcapillary. While certain types of chimerisms can be induced at different stages of generation (10, 11), it requires the use of expensive tools for manipulation and precision operators’ skills. In particular, as the success of the chimerism is influenced by the technique of the experimenter, the reproducibility of the experiment is considered difficult.

In the aggregation method, the embryos and stem cells are placed in small indentations on a petri dish and are allowed to remain adhered for inducing chimerism. This is a primitive method that can be used to produce chimeric embryos easily and inexpensively. Although several technical improvements have been proposed (12), the resulting chimeric embryos cannot be subjected to quantitative analysis as it is difficult to control the number of stem cells. In addition to the aforementioned injection and aggregation methods, a microaggregation method has also been developed (13), but no other methods are available for producing chimeric embryos.

To this end, this present study aimed to improve on the conventional methods for producing chimeric embryos. We attempted to apply the zona pellucida reconstruction method to the production of chimeric embryos. During the reconstruction of the zona pellucida, we attempted to isolate the cells within the oil droplets using microfluidic channels. Droplet microfluidics has been employed as an ultrahigh-throughput assay technology for a wide range of biological applications, including as antibody screening (14) and single-cell RNA sequencing (15). Water-in-oil (W/O) droplets formed by microfluidic devices are generally monodispersed, thus allowing the high-throughput creation of millions of tiny ‘test tubes’ that are represented by the individual droplets. Typically, the surfactants dissolved in the fluorinated oil stabilize and maintain the W/O droplets for over a month. In addition, the surfactants are non-toxic to mammalian cells, and the cells can therefore be cultured for up to two weeks (16). In summary, we attempted to produce chimeric embryos using the oil droplets and describe the novel method established in this present study.

## Materials and Methods

### Custom chips

The microfluidic droplet generator used in this study was designed on the CAD software Rhino 6 (McNeel & Associates, USA). A polymer mold was used for fabricating the droplet generator, which was prepared by using a 3D stereolithography tool (Acculas SI-C1000, D-MEC, JAPAN) that constructs three-dimensional structures layer-by-layer with epoxy-based UV curable resin (KC-1257, D-MEC) on a glass substrate. The 3D polymer mold was developed by using a solvent (EE-4210, Olympus, JAPAN) for at least 30 min to remove the uncured resin. The polymer mold was subsequently rinsed with ethanol and thoroughly dried on a hotplate at approximately 70°C, following which parylene C was vapor-deposited on the polymer mold using a parylene coating system (SCS Labcoter, USA). The droplet generator devices were fabricated with polydimethylsiloxane (PDMS, Sylgard 184, Toray Dow Corning). The microfluidic channels in the devices were filled with 2% trichloro silane (Sigma-Aldrich) in HFE7500 (3M, USA). The solution was removed and the devices were baked at 120°C, which increased the hydrophobicity of the microfluidic channels.

### Cells and embryonic culture

EGFP-expressing mouse ESCs, which have been established and described in the previous report (17), were cultured in ESGRO medium (Merck) supplemented with 20% KSR (Gibco).

All the protocols for the animal experiments were approved by the Animal Care and Use Committee of Kyushu University (protocol number: A30-304). All the mice were purchased from Japan SLC, Inc (Hamamatsu, Japan). The embryos were collected from superovulated B6D2F1 female mice at the 2-cell stage, at 1.5 dpc (days post coitum) and cultured in M16 medium in an atmosphere of 5% CO2 in air at 37 ºC.

### Formation of Chimeric embrys

After denuding the embryos at the morula stage with acidic Tyrode’s solution (Sigma) and EGFP-ESCs were transferred to a custom chip, the droplets were generated using a droplet generator (On-chip Biotechnologies Co., Ltd.) at the sample pressure and the oil pressure of 8.0 kPa. Fluorinated oil (008-FluoroSurfactant in HFE7500, RAN Biotech., Inc., MA, USA) was used in combination with surfactants at a concentration 2.06%. The resulting emulsion was collected in a 0.2 ml PCR tube, covered with PBS, and cultured in a CO_2_ incubator. After mixing the emulsion with an equal volume of 10% PFO (1H,1H,2H,2H-Perfluoro-1-octanol, Fujifilm, Osaka, Japan) in HFE7500, embryos were recovered from the droplets, washed with M2 medium, and subsequently cultured in M16 medium. Cell viability was measured by adding Propidium Iodide (PI) solution.

### Statistical analysis

Values of *P*<0.001 were considered to be statistically significant in the binominal tests.

## Results and Discussion

### Encapsulation of mouse ESCs in W/O droplets using microfluidic chips

The custom microfluidic chips were created by the method previously described in the Materials and Methods section (Fig. 1A). A highly magnified image of a microfluidic channel is depicted in Fig. 1B. The width of the channel in the microfluidic chip at the droplet generation point was 120 µm, and the flow of oil and cell suspension has been indicated in the figure by black arrows. The droplets were generated at the intersection of the oil channels and the cell suspension channel (Fig. 1C).

**Fig. 1.**
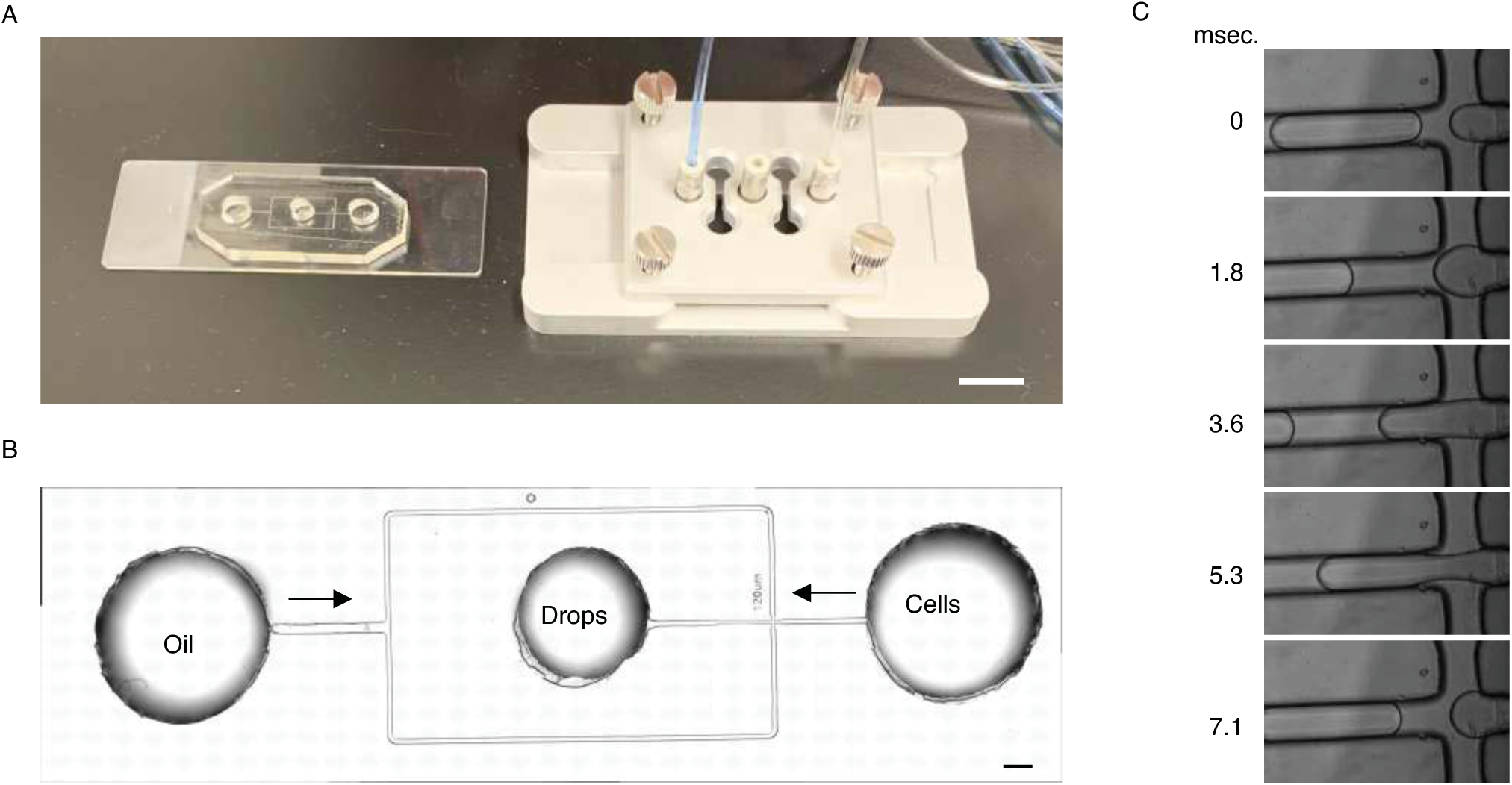
Custom chip and microfluidic channel. (A) Image of custom chip and pressure device; bar = 1 cm. (B) Magnified image of a microfluidic channel on custom chip. The black arrows indicated the direction of liquid flow; bar = 500 µm. (C) Time-scale representation of the droplets generated on the microfluidic channels.

EGFP-expressing mouse ESCs, that had been established and described in our previous study (17) were used. The ESCs expressed EGFP and showed naive-typed colony morphology (Fig. 2A). After pressurizing the cultured ESCs suspension with the oil in the custom microfluidic chips, the ESCs were successfully encapsulated and isolated into the droplets (Fig. 2B). In order to control the number of cells that were incorporated into each droplet, droplet generation was performed by altering the concentration of mouse ESCs in the suspension. A 2-fold dilution series of the suspension was prepared, ranging between 4.8 × 10^6^ cell/ml to 0.075 × 10^6^ cell/ml. The distribution of the number of cells isolated in the droplets using smear preparation of the generated emulsion is depicted in Fig. 2C. The results demonstrated that the cells could be isolated in half of the droplets at a concentration of 1.2 ×10^6^ cell/ml, and in 90% of the droplets at concentrations of 2.4 × 10^6^ cell/ml or higher.

**Fig. 2.**
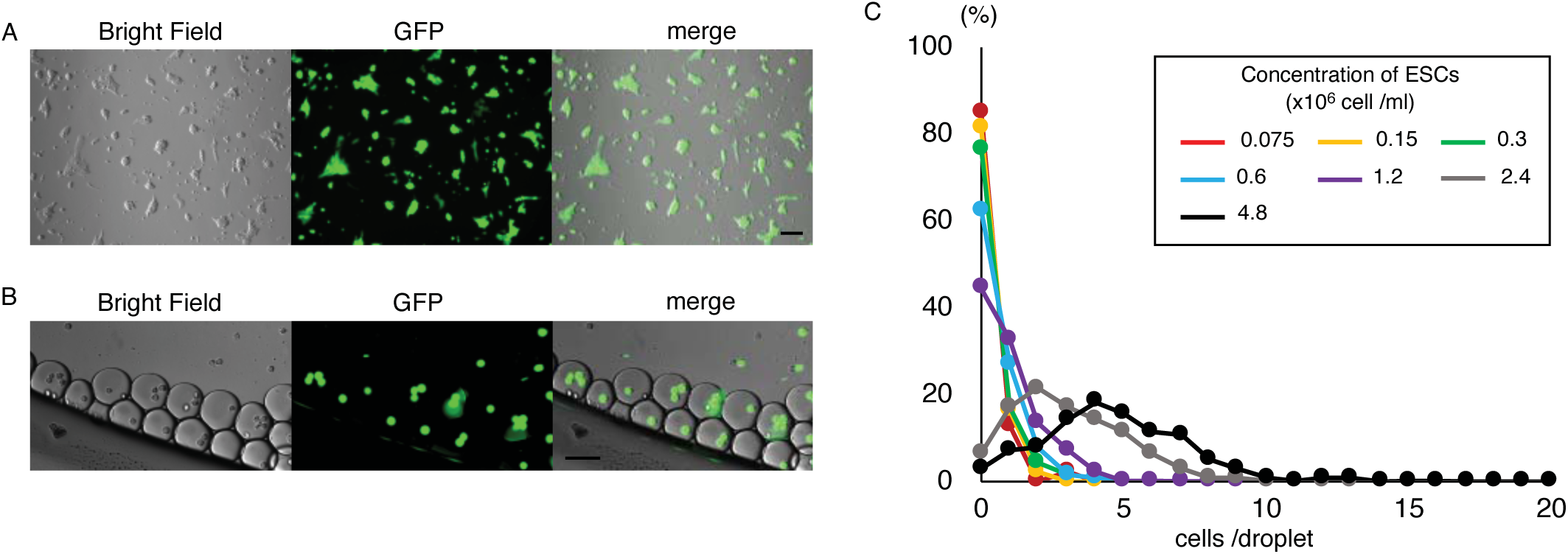
Isolation of mouse ESCs in the droplets. (A) Morphology and fluorescence imaging of EGFP-expressing mouse ESCs that were subsequently used for the formation of the chimeric embryos; bar = 100 µm. (B) Mouse ESCs isolated in the droplets. bar = 100 µm. (C) Distribution of the concentrations of mouse ESCs in the suspension and the number of cells encapsulated in the droplets.

### In-droplet culture of ESCs

In order to determine the period for which the ESCs could be cultured in the droplets, the survival rate following their in-droplet culture was measured. The generated droplets were cultured in 0.2 ml PCR tubes by overlaying with PBS to avoid evaporation of the volatile fluorinated oil (Fig. S1). After culturing the droplet-encapsulated ESCs for 1 and 2 days, it was observed that most of the cells became PI-positive and did not survive (Fig. S2). The cell viability was therefore subsequently measured 3 hourly for 12 hours, to reduce the culture period in the droplets to 1 day or less. It was observed that although the cell viability gradually decreased over the duration of the culturing (Fig. 3A), 75% of the ESCs survived for 9 hours and 80% survived for 6 hours (Fig. 3B). We therefore considered a culture time of 6 hours for ensuring a high survival rate in the subsequent experiments for producing chimeric embryos. Although the cause of cell death in the in-droplet culture after 1 day was unclear, it could be assumed that the ESCs did not survive due to cellular auxotrophy. As the auxotrophy of HEK cells and Jurkat cells are different, the survival rates of these immortalized cell lines in the in-droplet culture could have been different (16). Mouse ESCs are specifically auxotrophic for substances such as methionine (18), suggesting that cell death could have been induced by the low nutritive environment, due to their isolation into microdroplets. By modifying the continued culture of ESCs within the droplets, the method can be applied to the formation of embryoid bodies of uniform sizes (19), which might improve the quality of differentiation.

**Fig. 3.**
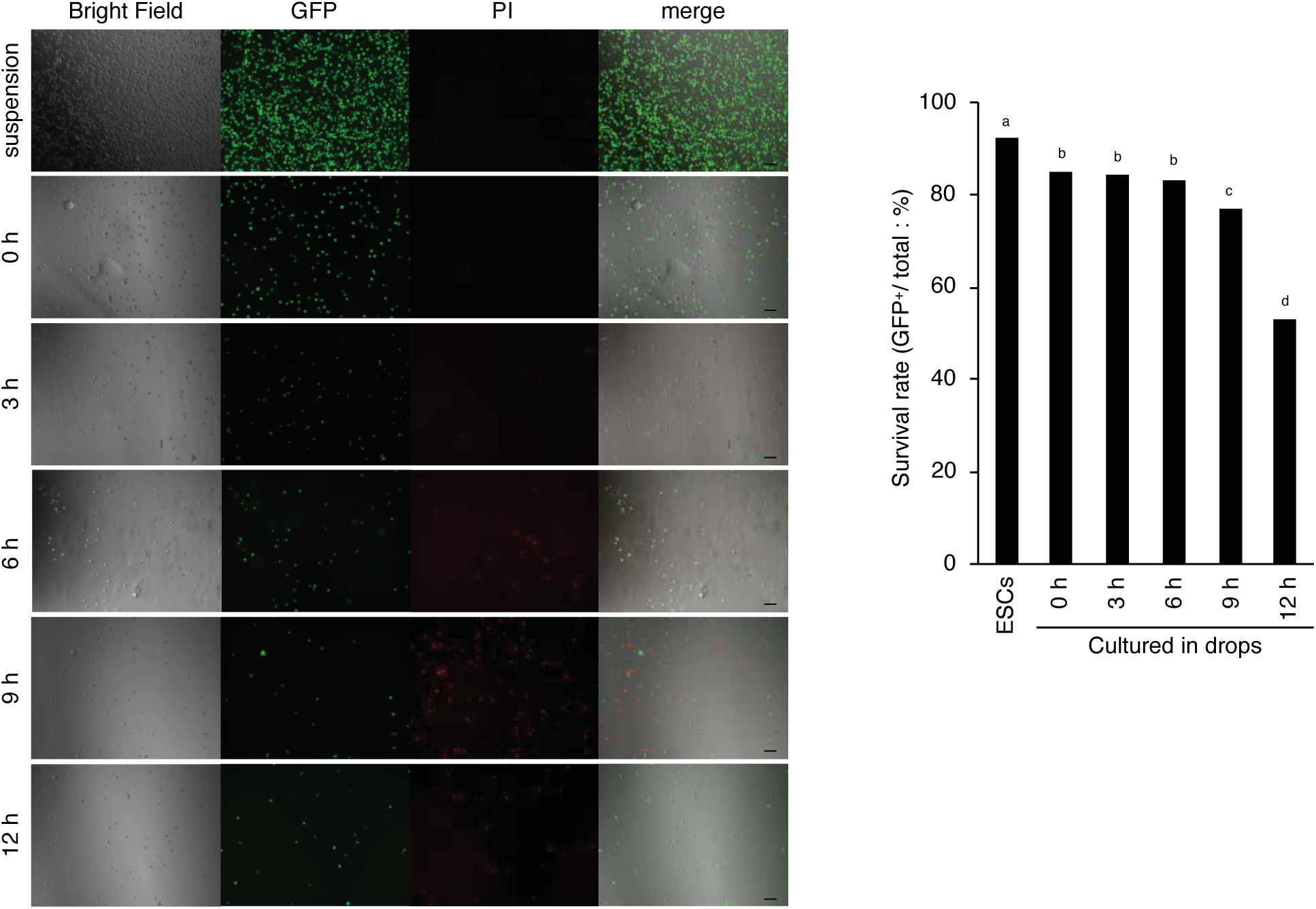
In-droplet culture of mouse ESCs. (A) PI-stained image of the EGFP-expressing mouse ESCs following in-droplet culture. The suspension prior to droplet generation was used as the control; bar = 100 µm. (B) Cell viability following in-droplet culturing. The number of PI positive cells were measured and represented the dead cells, while the EGFP-positive cells represented the living cells; a-d, indicates the significant differences at *P* <0.001.

### Production of chimeric embryos via droplets

A schematic diagram of the in-droplet cell culture experiment for the formation of the chimeric embryos is depicted in Fig. 4A. The details of the experimental procedures have been previously described in the Materials and Methods section. By adding the denuded embryos to the ESCs suspension and pressurizing with custom chips, we succeeded in isolating the embryos and ESCs into the same droplets (Fig. 4B). Following in-droplet culture, EGFP-positive cells were detected in morula stage (Fig. 4C). At 3.5 dpc, blastocysts with EGFP-positive cells in the inner cell mass were observed (Fig. 4D), indicating that the formation of chimeric embryos via the oil droplet method was successful. Table 1 enlists the production rate of chimeric embryos for the corresponding concentrations of ESC suspensions.

**Fig. 4.**
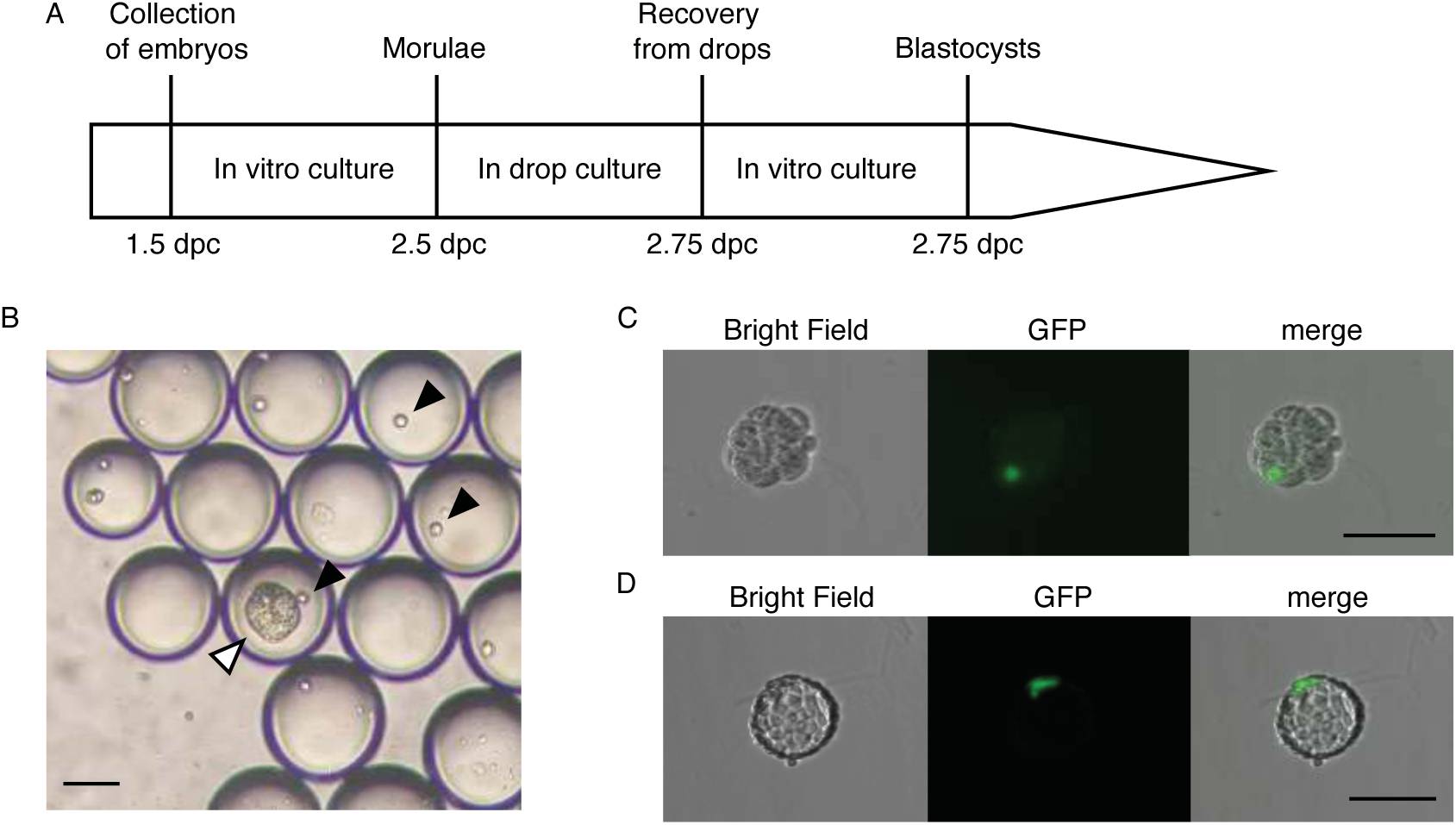
Formation of mouse chimeric embryos via the oil droplet method. (A) Schematic diagram of the experimental protocol. The morula stage embryos were co-cultured with EGFP-expressing mouse ESCs within the droplets for 6 hours. (B) The embryos at morula stage and the ESCs can be seen encapsulated in the droplets. The white arrowhead indicates the embryo at the morula stage, while the black arrowhead indicates the ESCs; bar = 100 µm. (C) A chimeric embryo recovered from the droplets at 2.75 dpc; bar = 100 µm. (D) Development of the recovered chimeric embryos in (C) into blastocyst stage; bar = 100 µm.

**Table 1.** Production efficiency of chimeric embryos

| Concentration<br>of ESCs (cell/ml) | Total number<br>of embryos | The number of<br>chimeric blastocysts | chimerism<br>(%) |
| --- | --- | --- | --- |
| 1.2 x 10<sup>6</sup> | 34 | 9 | 26<sup>a</sup> |
| 2.4 x 10<sup>6</sup> | 64 | 44 | 69<sup>b</sup> |
| 4.8 x 10<sup>6</sup> | 93 | 82 | 88<sup>c</sup> |

There are two methods for reconstructing the embryonic zona pellucida, one using agarose capsules (20) and other using sodium alginate capsules (21). Both methods have disadvantages, in that they have complicated process of production or inhibition of embryonic development (22, 23). In this present study, it was observed that a 6-hour embryonic in-droplet culture produced no effects on the subsequent development of the embryos. Additionally, the embryos could be easily recovered from the droplets by simply by adding PFO solution.

In this present study, we established the droplet method for producing chimeric embryos, which is completely different from the conventional injection and aggregation methods that are used for the generation of chimeric embryos. This method allows a superior for the production of chimeric embryos, thus contributing to the efficient production of chimeric animals in the future.

## Acknowledgements

This work was supported by the 40th LNest Grant and the Qdai-jump Research (QR) Program. The authors are grateful to Dr. Kiyoshi Kano of Yamaguchi University for providing the mouse ESCs in accordance with Research Results Materials Handling Regulations of Yamaguchi University.

---

**Fig. S1.**
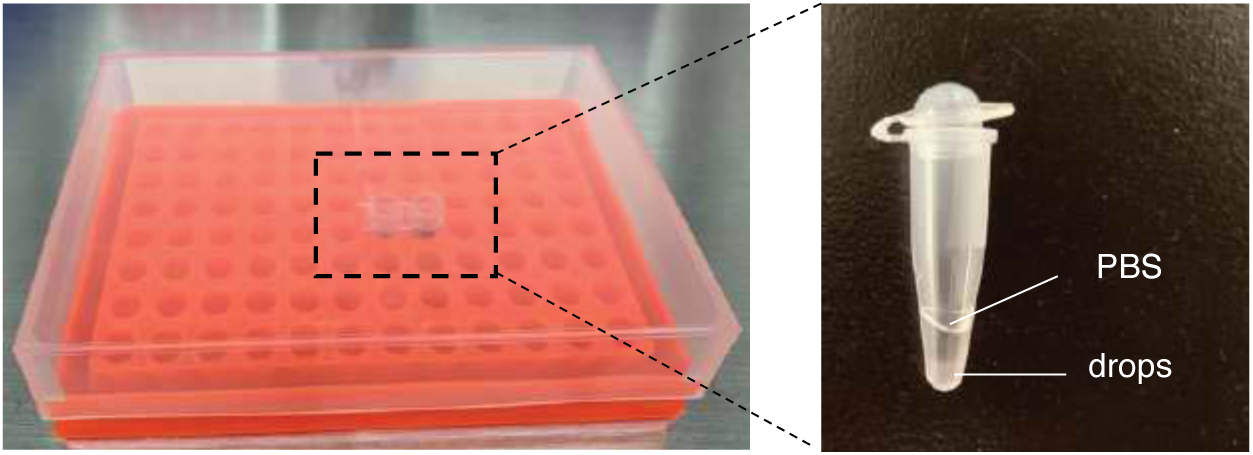
The in-droplet culture method. The droplets were layered on PBS and cultured in PCR tubes (left) with caps half open or fully open on PCR racks (right).

**Fig. S2.**
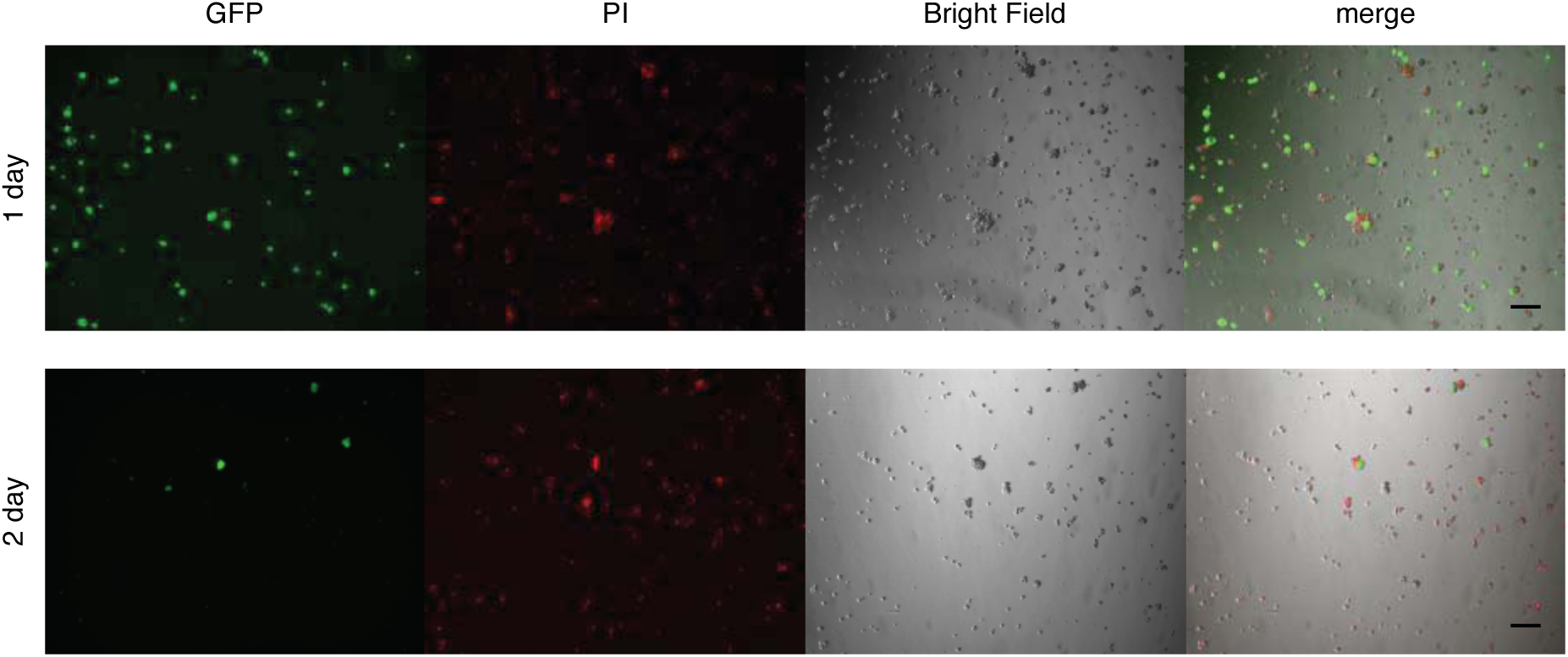
In-droplet culture of mouse ESCs for 1-2 days. Fluorescent imaging using PI and EGFP after 1-2 days of in-droplet culture. Most of the cells were founded to be PI positive; bar = 100 µm.

